# Reconstructing wells from high density regions extracted from super-resolution single particle trajectories

**DOI:** 10.1101/642744

**Authors:** P. Parutto, J. Heck, M. Heine, D. Holcman

## Abstract

Large amount of super-resolution single particle trajectories has revealed that the cellular environment is enriched in heterogenous regions of high density, which remain unexplained. The biophysical properties of these regions are characterized by a drift and their extension (a basin of attraction) that can be estimated from an ensemble of trajectories. We develop here two statistical methods to recover the dynamics and local potential wells (field of force and boundary) using as a model a truncated Ornstein-Ulhenbeck process. The first method uses the empirical distribution of points, which differs inside and outside the potential well, while the second focuses on recovering the drift field. Finally, we apply these two methods to voltage-gated calcium channels and phospholipids moving on the surface of neuronal cells and recover the energy and size of these high-density regions with nanometer precision.

## 1 Introduction

Super-resolution Single Particle Trajectories (SPTs) are used to monitor the dynamics of large amount of particles that can be cytoplasmic molecules, membrane receptors or channels in live cells. Over the past decade, statistical methods based on stochastic models have been developed to segment [1, 2], to interpret these large data sets and to extract relevant biophysical parameters such as flows, fluxes and even statistics of arrival times between various subregions [3, 4, 5, 6, 7, 8]. The most striking and universal characteristic of these trajectories is that they are not homogeneously distributed in cells, but rather are concentrated in sub-regions, a phenomenon that is not fully understood: what are these regions of high densities? What are the underlying physical forces that restrict and confine the trajectories? For example, AMPA receptors that traffic on the surface of neuronal cells accumulate specifically at the post-synaptic density (PSD) of synapses, where they are needed for proper synaptic transmission [9, 10]. Similarly, at the pre-synaptic terminal, voltage-gated calcium channels (CaV) can accumulate on membrane subregions, with a size around hundreds of nanometers [11]. Retaining these channels guarantee that calcium ions can flow near vesicles to trigger release.

Heterogeneous distribution of trajectories within specific regions of large density were observed systematically for a large spectrum of membrane proteins, channel receptors [3, 11]: they are often concentrated in specific regions for a physiological purpose, however, the biophysical mechanism for such accumulation remains unknown. A possible mechanism to retain trajectories is a converging field of force due to the presence of an extended potential well. These structures have been detected with a size of hundreds of nanometers [3, 12, 11]. However the physical origin of these wells remains unclear because the length of classical electrostatic interactions is ten times shorter [13]. These high density regions are characterized by several features: 1) a converging drift field whether or not it is the gradient of a potential energy, 2) an energy depth and 3) a boundary. Finding and estimating these geometrical characteristics from single trajectories and their statistical distribution remain challenging especially at tens of nanometers below the diffraction limit.

Here, we present two statistical methods to detect and interpret highdensity regions contained in single particle trajectories. We reconstruct the possible underlying potential wells. The first approach is based on estimating the density of points of a truncated Ornstein-Ulhenbeck process (which account for a motion driven by a converging force and diffusion) as a stochastic model. We recover the center, the covariance matrix and the boundary of the wells. The second approach is based on estimating the drift flow. Both approaches are validated on stochastic simulations. Finally, we apply the present methods to estimate the characteristics of potential wells using SPTs of voltage-gated calcium channels (CAV) and phospholipids (GPI-GFP or lipid anchored GFP) embedded in the membrane of neuronal cells.

## 2 Methods

### 2.1 Coarse-grained description of stochastic trajectories

The Langevin’s description of a stochastic trajectory is summarized by the equation

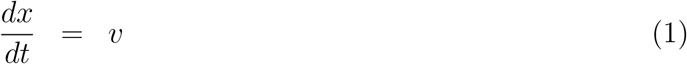

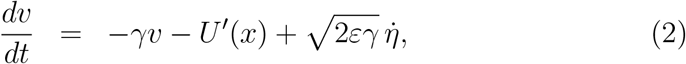

where *γ* = Γ*/m* is the dynamical friction coefficient per unit mass and U is a potential field. In the Smoluchowski’s limit [14, 15], the description reduces to

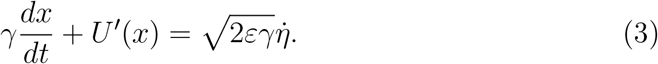

However, this model assumes that the diffusion of a protein or a particle embedded in a membrane surface is generated by a diffusion coefficient *D* and a field of force *F* (*X, t*),

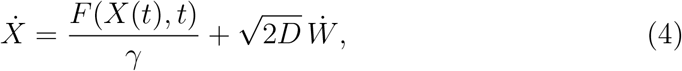

where *W* is a Gaussian white noise and *γ* is a uniform viscosity [14]. The source of the driving noise is the thermal agitation of the ambient lipid and membrane molecules. However, due to the acquisition timescale of empirical recorded trajectories, which is too low to follow the thermal fluctuations, rapid events are not resolved in the data, and at this coarser spatiotemporal scale, the motion is described by an effective stochastic equation [3, 16]

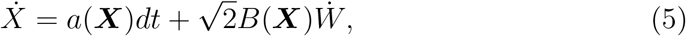

where *a*(***X***) is the drift field and *B*(***X***) the diffusion matrix. The effective diffusion tensor is given by 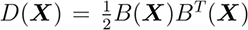 (*.^T^* denotes the transposition) [17, 14]. The diffusion tensor accounts for impenetrable obstacles of various sizes. Note that the interpretation at the physical level of the stochastic equation is from the Ito’s sense and not Stratanovich or any other sense, because a physical process has to be non-anticipating [17] (the future cannot interfere with the past).

### 2.2 Potential well characteristics

The drift field *a*(***X***) in equation 5 may represent a field force that acts on the diffusing particle, that could due to a potential well [13]. When the diffusion tensor ***D***(***X***) is locally constant and the coarse-grained drift field *a*(***X***) is a gradient of a potential

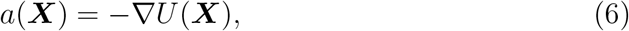

then the density of particles is given locally by the Boltzmann distribution [18]

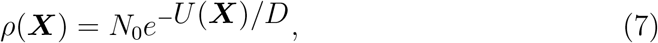

where *N*_0_ is a normalization constant. An infinite paraboloid potential well with an elliptic base has the analytical representation for ***X*** = (*x, y*)

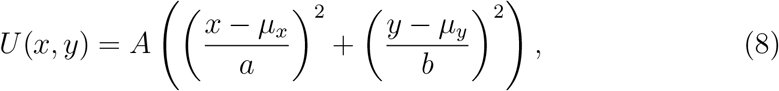

where the center is (*µ*_*x*_, *µ*_*y*_) with amplitude *A* and *a, b* are the sizes of the semiaxes of the ellipse. To account for a finite well, we restricted the influence of the well to the region

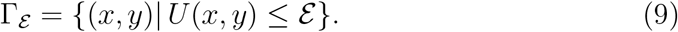

The truncated energy function *U* associated to a parabolic potential well is finally given by

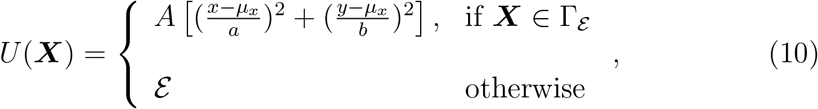

from which the drift field is the gradient of the energy, given by

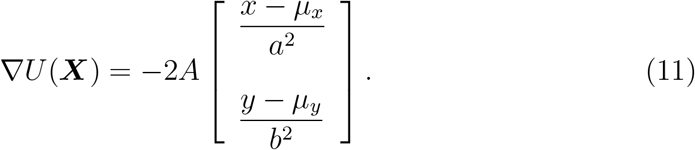

The goal of the present work is to recover, from empirical single particle trajectories that consists of few successive points acquired at a sampling rate ∆*t*, the center (*µ*_*x*_, *µ*_*y*_), the amplitude *A*, the size of each semi-axis *a, b* and the boundary *ε*.

### 2.3 Simulations of stochastic trajectories

To validate the two methods, we first generated synthetic single particle trajectories from the stochastic process

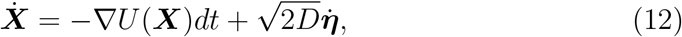

using the classical Euler’s scheme. The potential *U* is defined in equation (8), *D* is the diffusion coefficient and ***η*** is the normalize Gaussian noise. In practice, we follow the experimental protocol and generated at a time resolution ∆*t N* trajectories (*X*_1_(0),.., *X*_*N*_ (*K*∆*t*)) containing *K* points (*K* = 20), summarized in Fig. 1A-B).

We consider two types of numerical simulations depending whether the initial points *X*_*i*_(0) are uniformly distributed 1) inside the well or 2) inside a square box surrounding the well. This uniform distribution represents the random activation of fluorophores by a laser (Fig. 1C). To guarantee a constant number of points inside a well across various simulations, we generated new trajectories from an initial uniform distribution until we reached a desired number of points inside the well. This resetting procedure generates a distribution of points which depends on the initial uniform distribution. However, in the limit of large *N*, the distribution of points converges toward the steady-state, which is Gaussian inside the well and uniform outside, when trajectories are confined to large square domain.

**Figure 1:**
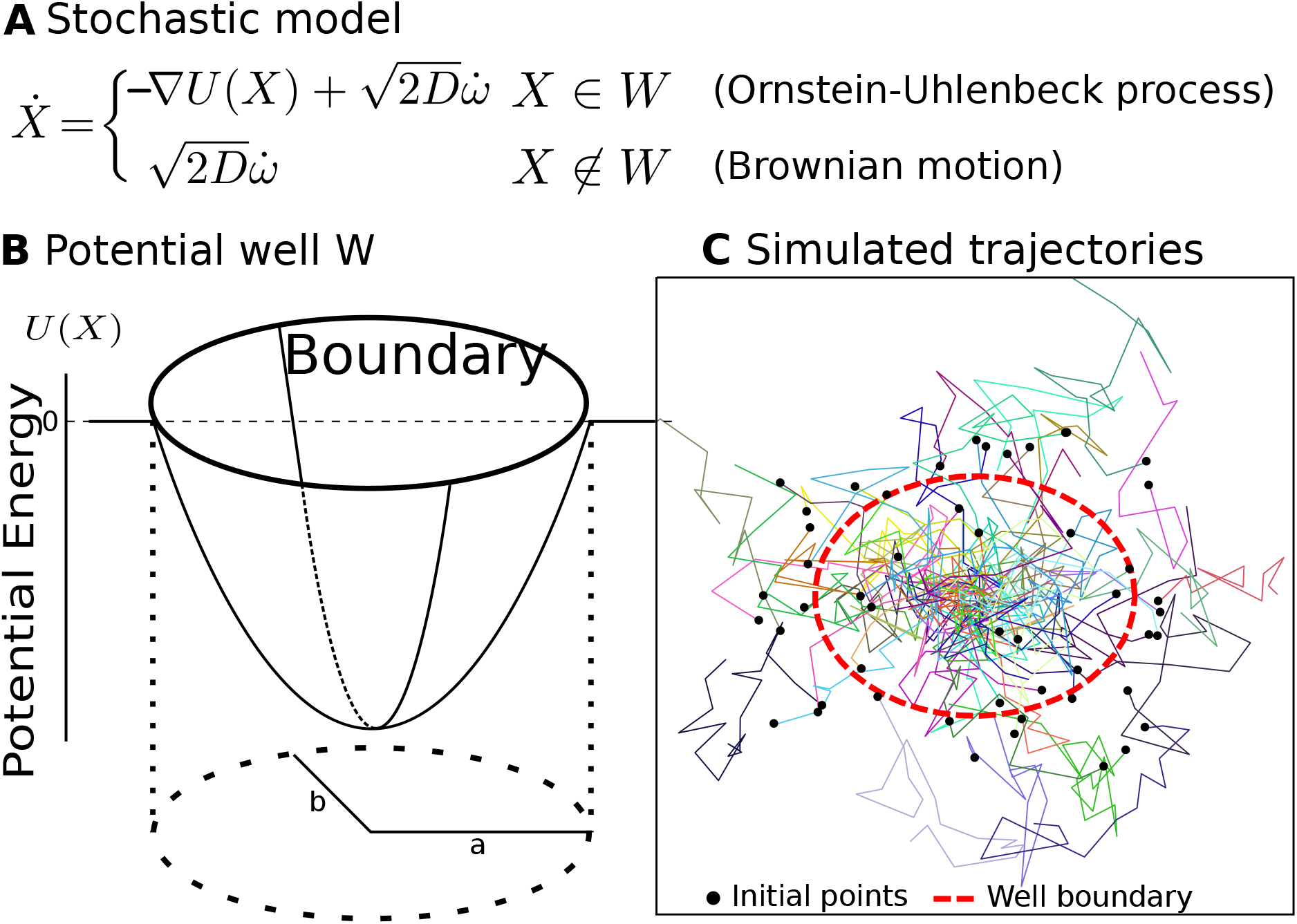
Stochastic modeling and numerical simulation scheme: **A** Ornstein-Uhlenbeck stochastic process truncated to well and Brownian outside used to generate single trajectories. **B** Model of a truncated potential well with two axes *a, b* and energy *U* (*X*) with a boundary. **C** Trajectories generated with equation presented in **A**. The initial points (black dots) can either be located inside or outside the boundary of the well (dashed red). Parameters: *D* = 0.042*µm*^2^*/s*, *λ*_*x*_ = 10, *λ*_*y*_ = 17.78.

### 2.4 Empirical estimators

The drift of the stochastic model 5 can be recovered from SPTs acquired at any infinitesimal time step ∆*t* by estimating the conditional moments of the trajectory increments ∆*X* = *X*(*t* + ∆*t*) *− X*(*t*) [14, 19, 20, 16, 21]

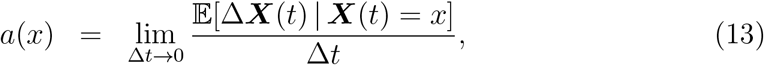

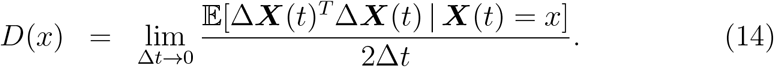

The notation 𝔼[.|*X*(*t*) = *x*] means averaging over all trajectories that are at point ***X*** at time *t*. To estimate the local drift *a*(***X***) and diffusion coefficients *D*(***X***) at each point ***X*** at a fixed time resolution ∆*t*, we use a similar procedure as the one for the estimation of the density in section 3 based on a square grid. The points of trajectories are first grouped within small square bins *S*(*x*_*k*_, *r*) of size ∆*x* and centered on lattice grid *x*_*k*_ and the drift and local diffusion coefficient are estimated for each of the square.

When there are *N* trajectories {*X*_*i*_(0)*, …, X_i_*(*K*∆*t*)}, with *i* = 1 … *N* and ∆*t* the sampling time, the discretization of equation 13 for the drift *a*(*x*_*k*_) = (*a*_*x*_(*x*_*k*_), *a*_*y*_(*x*_*k*_)) in a bin centered at position *x*_*k*_ is

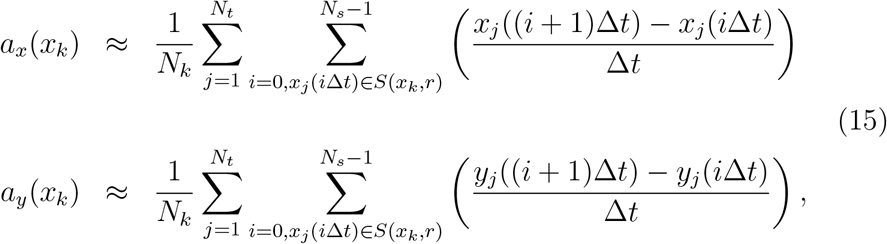

where *N*_*k*_ are the number of points of the trajectory falling in the square *S* (*x*_*k*_, *r*). Similarly, the components of the effective diffusion tensor *D*(*x*_*k*_) are approximated by the empirical sums

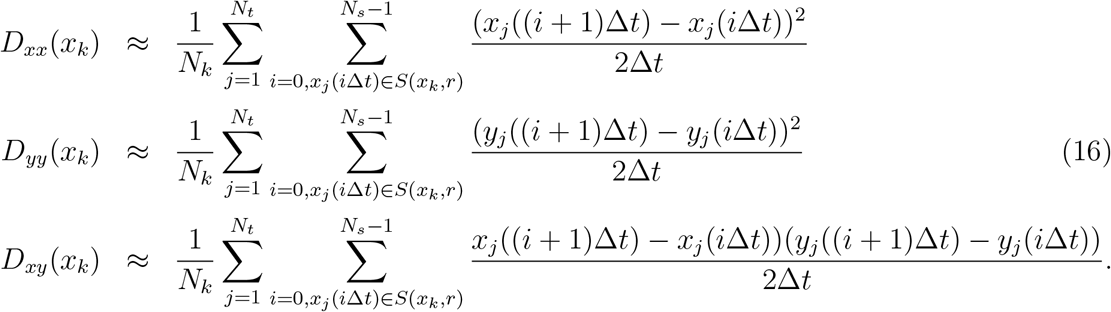

There are thus several free parameters such as the position of the grid, the bin size ∆*x* that should be fixed and optimized during the estimation procedure.

### 2.5 Correcting the drift estimation

We recall briefly here (see SI) a correction term to be added in order to recover Ornstein-Uhlenbeck process of parameter *λ* and centered at *µ* (eq. (12)). We derived in the SI that the drift term at position *x* and at resolution ∆*t* is given by

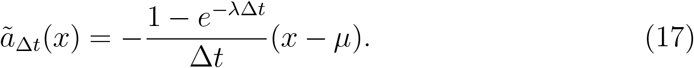

Hence, the first order moment at resolution ∆*t* computed from the displacement *X*(*t* + ∆*t*) − *X*(*t*) from the empirical data deviates from the expected drift. When *λ*∆*t* is small, the first order Taylor expansion leads to the approximation

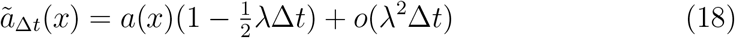

and hence to recover the drift, we have to account for the factor 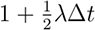 on the estimated drift.

### 2.6 Data processing

For the experimental data related to CaV2.2 data, we refer to [22], while the GPI-GFP data have been described in [23]. Here, we used the following procedure to process both CaV and GPI datasets: to remove any possible motion of the wells due to cell motion or drift acquisition that would affect the analysis of the trajectories, we initially isolated trajectories in non-overlapping time windows of 20s. For each time window, we use a square grid with bin sizes ∆*x* around trajectories and isolated the 5% highest density bins as possible regions containing potential wells. For each selected bin, we detected well as follows: we first used 90% of the local density (threshold *α* = 0.1) to detect the center of the well from equation (21).

Using this initial possible point for the center, we computed by iteration, the annulus starting at the radius *r*_*min*_ and ending at *r*_*max*_, with a width increment ∆*r*. We estimated the covariance ratio (eq. (24)) and density from each annulus (*r, r*+∆*r*). To obtain the semi-axes ratio, we toke the maximum value of the covariance ratio until a maximum distance *r*_*cov*_ = 150nm. In that case, the radius *r*_0_ is obtained by using the density curve at the level line *T_ρ_* = 35%. Once the center and semi-axes of the well are found, the diffusion coefficient is determined using eq. (16) using all displacements where the initial points fall inside the well.

## Results

### 3 Recovering a bounded potential well from the point density of trajectories

We reconstruct the characteristics of the well from the distribution of points resulting from SPTs. This approach ignores the time series of the trajectories and relies on the a priori knowledge that the well is a truncated paraboloid. To this end, we recover the center and covariance matrix of the steady-state density distribution using the binning of trajectory points into a square grid (Fig. 2A). We recall that the steady-state is the Boltzmann distribution, which is the solution the Fokker-Planck equation 8: it is given inside the well by

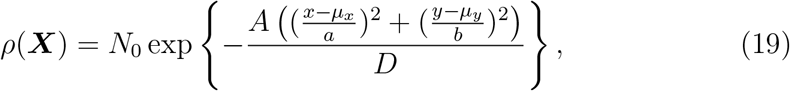

where *N*_0_ is a normalization coefficient (the other parameters are defined in subsection 2.2). To select parts of trajectories inside the well, we use the ensemble

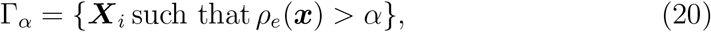

where *ρ*_*e*_ is the empirical point density estimated over the bins of the square grid constructed from the ensemble of trajectories (Fig. 2B). The ensemble Γ_*α*_contains all trajectory points falling into a bin with a density greater than the density threshold *α*.

**Figure 2:**
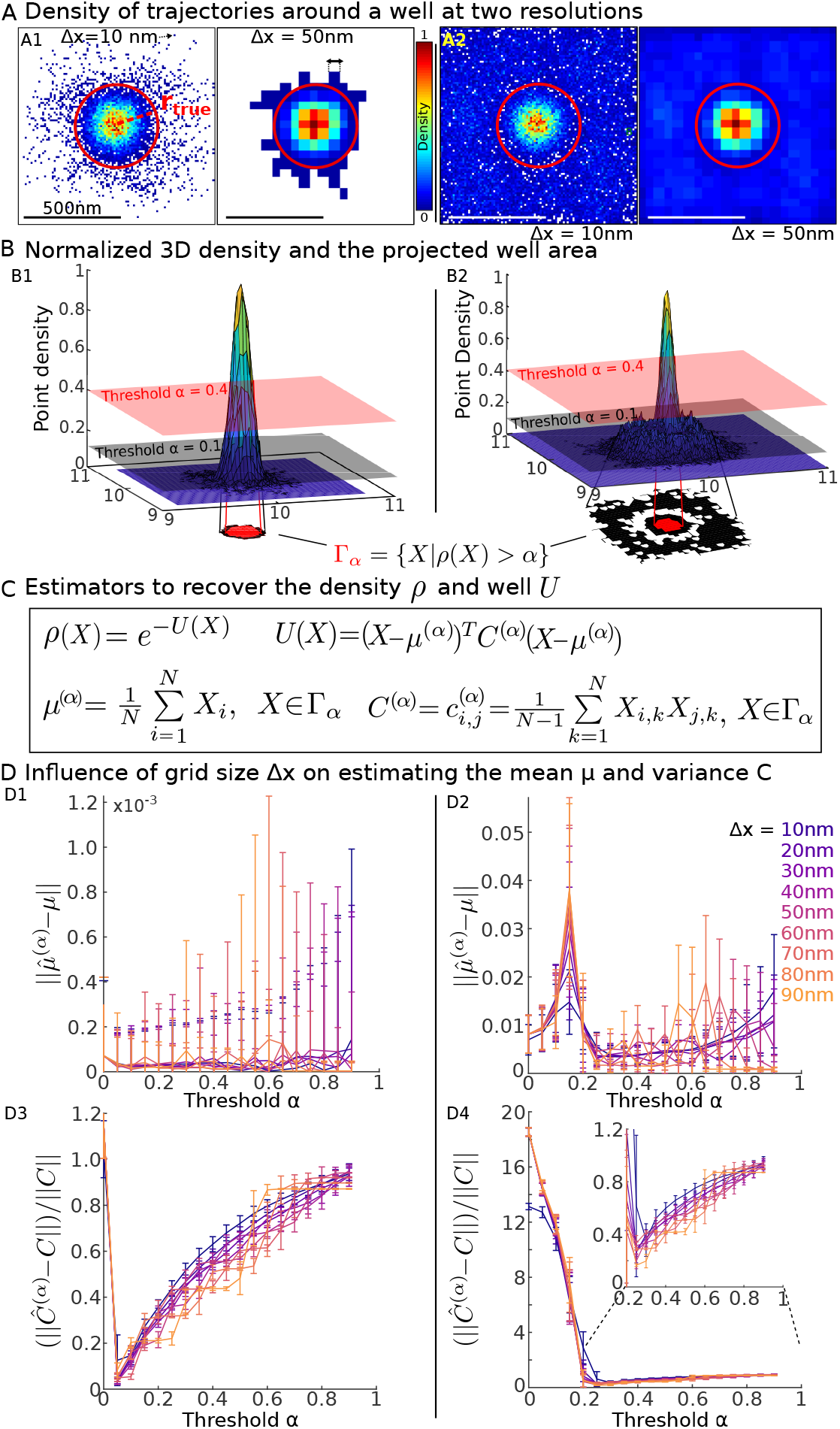
Recovering a truncated potential well from the density of points. **A** Density maps (in log(points)/*µm*^2^) for two different grid sizes ∆*x* = 10 (left) and 50*nm* (right) when the initial points are located inside the well (A1) or uniformly distributed in a square of size 1*µm* (A2). **B** Normalized three-dimensional empirical density function *ρ* obtained from (A). We plotted the ensemble Γ_*α*_= {*X |ρ*(*X*) *> α*} for *α* = 0.1 (black) and *α* = 0.4 (pink) and the projected area (red) in the well in the two cases (B1) and (B2) associated to (A1) and (A2) respectively. **C** Empirical estimators used to recover the characteristics of the well *U*: the center of the well and the covariance matrix are estimated by *µ*^(*α*)^ and *C*^(*α*)^ respectively, which are the average of the points and the quadratic sum, restricted to the ensemble Γ_*α*_. **D:** Influence of the grid size ∆*x* and threshold *α* on the well characteristics estimations. D1,D3 (resp. D2, D4) panels are obtained by computing with the initial distribution described in A1 (resp. A2).

### 3.0.1 Estimating the center (*µ*_*x*_, *µ*_*y*_) of the well

To recover the center of the distribution, we consider all points ***X***_*i*_ = (*x*_*i*_, *y*_*i*_) located in Γ_*α*_ (see (20) and Fig. 2C) and use the empirical estimators:

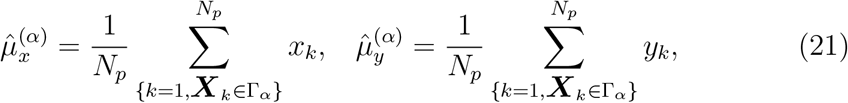

where *N*_*p*_ is the number of points in the ensemble Γ_*α*_.

### 3.0.2 Estimating the covariance matrix of data

To estimate the covariance two-by-two matrix *C*^(*α*)^, we start from the representation

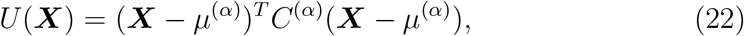

and use the empirical estimators

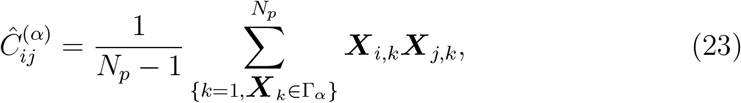

where ***X***_*i,k*_ is the *i*^*th*^ coordinates of ***X***_*k*_ (Fig. 2C). The accuracy of estimators (21) and (23) are analyzed by plotting the errors between the true and the estimated centers 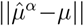 and the covariance matrices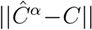 (*L*^2^ norm of the matrix) versus the parameter *α*. We used various grid sizes, varying from ∆*x* = 10 to 90*nm*: when *α* decreases from one to zero, when the initial points are located inside the well, the position of the estimated center converges toward the true one and the fluctuations (SD computed over 100 realizations) are decreasing (Fig. 2D1-D2) with *α*. However, when the initial points were chosen also outside the well, we found that there was an optimal threshold value for *α ≈* 0.3 for which the error in the center positions is minimum. Below this value, the points falling outside the well are also contained in Γ_*α*_, thus contaminating the estimation. When the initial points only fall inside the well (Fig. 2D3), the ensemble Γ_0_ contains external trajectories that affect the estimation of covariance matrix *C*^(*α*)^. As *α* increases, these external points disappear from Γ_*α*_and the error becomes minimal at the value *α_opt_* = 0.05. When *α* continues to increase, the estimators become less accurate. However, when the initial points were chosen also outside the well, the error starts by decreasing because the computation accounts for trajectories that are not inside the well (Fig. 2D4). As *α* increases, the estimator converges toward an optimal value *α* 0.25, which minimizes the error (that is 75% of the points are used). When *α* continues to increase, the error increases slowly, similar to the case of Fig. 2D3.

To conclude, having trajectories mainly inside or also outside the high density regions leads to different types of results. Moreover, while is it interesting to use as many points as necessary to estimate the center, to estimate the covariance matrix, we needed to truncate the density of points and found an optimal value for *α*, in order to reject the points that do no fall inside the well.

### 3.0.3 Estimating the boundary of the well

None of the estimators described in the previous subsections could be used to reconstruct the boundary of the potential well, thus we started with the recovery of a circular boundary and later on expand to an elliptic one in two cases: when the initial points falls only inside the well and when they can also fall outside.

The first step consists in discriminating the nature of the boundary between a circular and elliptical. To do so, we computed from the matrix (23), the covariance ratio

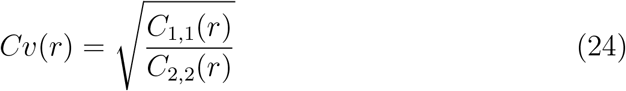

estimated over the empirical data inside the annulus (*r, r* + ∆*r*). We recall that the diagonal form can be found from equations (19) and (22), where the covariance matrix is given by

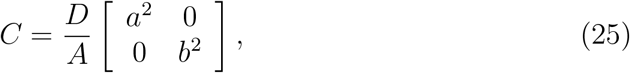

and thus in that case, we expect that the covariance ratio is 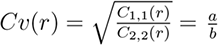: it is precisely equal to the semi-axes ratio and does not depend on any other parameters. In the case of a disk, the ratio is *Cv*(*r*) = 1, as shown in the simulation cases (Fig. 3A1-A2, A3-A4). Note that when the covariance matrix is recovered, the coefficients *λ*_*x*_ and *λ*_*y*_ can be recovered using equation (25) from the diffusion coefficient *D* and the semi-axes *a, b*

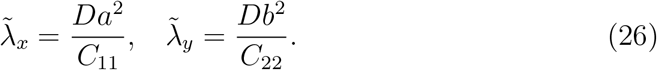

Once the well boundary has been identified as circular, to estimate the radius of the boundary *r*_0_, we plotted the density of points *ρ*(*r*) versus *r*, the radial distance with respect to the center 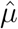 (estimated in subsection 3.0.1).

**Figure 3:**
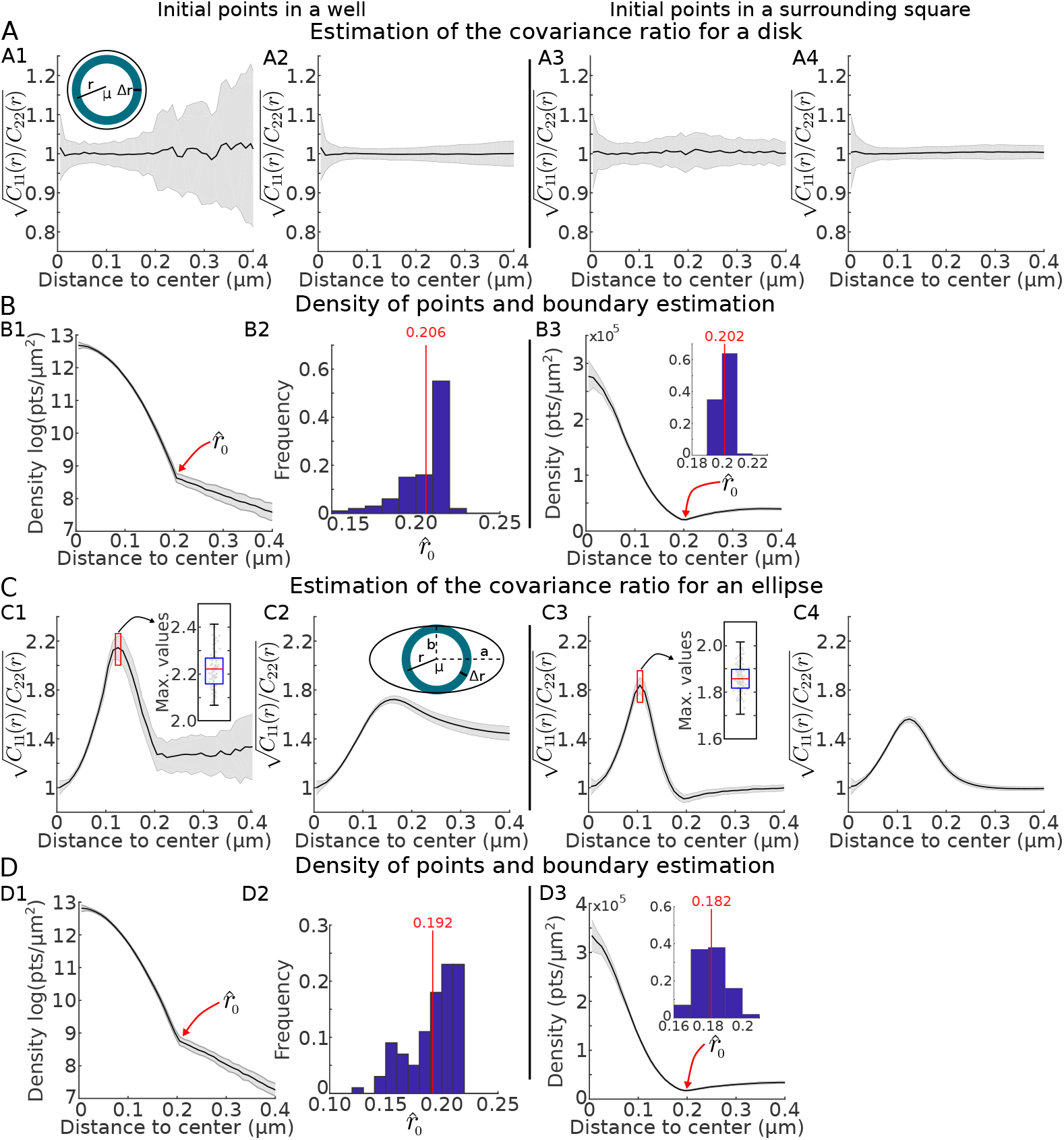
Estimating the potential well boundary. **A.** Covariance ratio 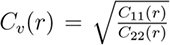 estimated in the annulus *r, r* + ∆*r*. Panel A1 (resp. A3) are for initial points distributed in a square (target disk resp). Panel A2 and A4 show the cumulative 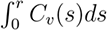.**B.** Point density estimated from the distance *r* to center. B1: Log of the density, showing a clear inflection point at the boundary of the well (criteria of selection) when the initial points of trajectories falls inside the circular well. B2: Estimation of the radius 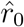 using a threshold at 4% of the total mass. B3: density of points obtained when the initial points start inside a square. The minimum is achieved near the boundary where we estimated radius 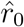 of the circular well. **C-D**. Generalization to the case of an ellipse of panels A-B. C1: the covariance ratio has a maximum when the moving disk of radius *r* becomes tangent to the smallest axis (inset boxplot for 100 simulations), from which we estimated the radius *b*. C2 cumulative. C3,C4 same as C1, C2 but for initial points inside the ellipse. **D.** Density of points is estimated inside the annulus using a corrected distance 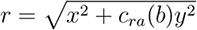 using the width of the elliptic base, estimated in panel C. The panel D1,D2 and D3 are equivalent to B1,B2 and B3.

Interestingly, this procedure emphasizes a local minimum, due to the difference between the Boltzmann distribution (attraction of the well) and the uniform distribution of the trajectories outside the well (Brownian motion). This difference in the distributions allows estimating 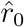 for the well radius *r*_0_ (Fig. 3B1). When the initial points falls inside the well, the density of points decays with the radius *r* and the boundary can be identified by plotting −log *ρ*(*r*). Indeed, for points falling inside the well, we have the approximation log *ρ*(*r*) ∼ *C*_0_ − (*αx*^2^ + *βy*^2^), where *r*^2^ = *x*^2^ + *y*^2^, with *α* = 2*A/a*^2^, *β* = 2*A/b*^2^ and exp *C*_0_ is the maximum value of the distribution. Outside the boundary, the distribution is usually sparse, generated by the Brownian motion. When trajectories are confined to a square, the empirical distribution outside the well converges to the uniform distribution as the number of points increases. When there are no confinement mechanisms, but trajectories have a finite length *K*, the distribution *q*_*K*_(***x***) of *K* successive points of a Brownian motion starting at uniformly distributed outside the well *S − C* is the sum

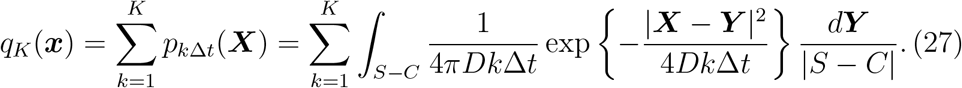

In the limit ∆*t* small, the Ornstein-Ulhenbeck and the uniform distributions are continuous at the boundary of the well, but the derivative is discontinuous. In practice, for the case of a circle, we can find the boundary by fitting a parabola *C*_0_ *− C*_1_*r*^2^ (with parameters *C*_0_ and *C*_1_) to *−* log *ρ*(*r*)) starting from zero and estimate the first point *r*_0_ of deviation of the error 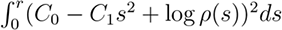.

In the case of an elliptic well, we modified the previous method as follows. The covariance ratio *Cv*(*r*) reaches a maximum when the radius *r* is equal to the smallest axis *b* of the ellipse (Fig. 3C, for a true ratio *a/b* = 2). After we found the maximum value for the ratio and estimated the length *r* = *b*, we constructed the elliptic distance 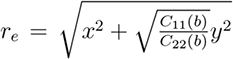, for any point *P* = (*x, y*). We then plotted the distribution of points according to the distance *r*_*e*_ (Fig. 3D). This procedure allows us to estimate the second elliptic semi-axis using the same procedure we have used for the disk (Fig. 3D1-D3).

To conclude, the present method based on density of points allows to reconstruct a finite parabolic potential well (center, boundary, small and large axes) with an elliptic base from the empirical density of points. In SI Figs. S1 and S2, we compare this density method with the classical MLE, which is used to recover the center and the axes, but not the boundary.

## 4 Estimating the characteristics of the well using the velocity distribution

In this section, we present a second method based on displacements to recover the drift of the vector field and reconstruct the following parameters of the potential wells: center *µ*, the two axes *a, b* and the boundary of the well. This method is based on least square quadratic error (LSQE), where we optimize the error

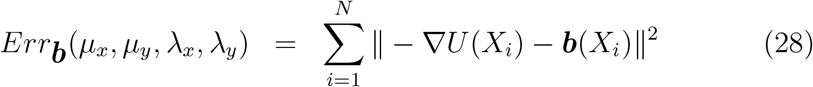

between the empirical drift ***b*** and the parabolic well *U*.

### 4.1 Estimating the center and the field coefficients of the potential well

By minimizing the error 28, we obtain the estimators for the center

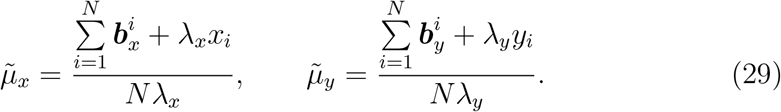

and for the eigenvalues of the covaraince matrix:

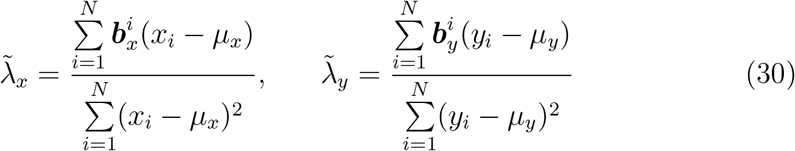

Combining equations 30 and 29 leads to an explicit expression of each parameter *µ* and *λ* (see SI). In Fig. 4A, we compare the reconstructed and the true drift based on equation (15) for various grid sizes. At this stage, we estimated the drift for bins that are falling inside the well (we assumed here that the boundary was known). The error of the norm 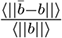 is plotted for multiple time steps ∆*t* and for three grid sizes ∆*x* = 10; 50 and 90nm in Fig. 4B. Having both a small grid size and time step ∆*t* produces a large error that quickly decreases with increasing the time step. Interestingly, for a large grid size, we found a slow increase of the error when increasing ∆*t*. To better understand which parts of the field contribute to the error, we plotted the error versus the distance to the center in Fig. 4C, showing that for small size ∆*x* = 10*nm*, a contribution comes from the center, while for large step ∆*x* = 50, 90*nm*, an error can come also from the estimations at the boundary.

**Figure 4:**
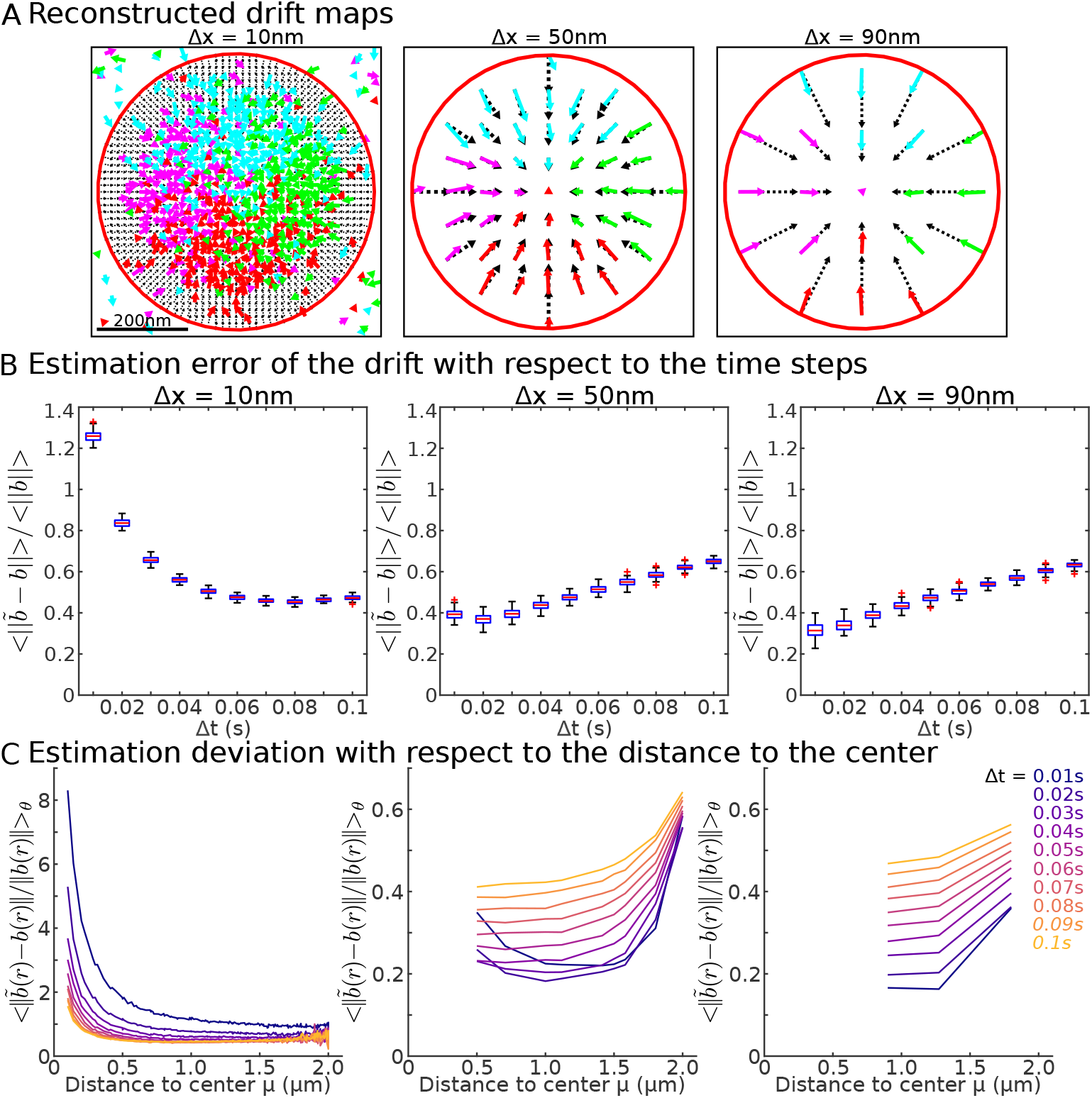
vector field characteristics. **A** Recovering the local drift field inside a circular well for different grid sizes (10*nm*, 50*nm*, 90*nm*) using numerical simulations with ∆*t* = 20*ms*, with the constraints that at leat 10 points falls inside a bin. **B** Error between the true and observed fields averaged over all the square bins inside the well vs the time step ∆*t*. **C** Error between the true and observed fields averaged over the radial angle vs the distance r to the center of the well center for various timestep (see color code).

Finally, to estimate the boundary of the well from the drift distribution (Fig. 5A), we plotted the drift amplitude as a function of the distance to the well center (Fig. 5B, blue cross representing the drift amplitude in one bin at distance r). As expected, from the distribution and the average (Fig. 5B lower panel), the boundary can be recovered from the local maximum: indeed, after the boundary is past, the contribution of the deterministic field disappears and it only remains in the statistics, the fluctuations due to the Brownian nature of the motion. We apply the same procedure for the case of an ellipse (Fig. 5C-D) and recover the boundary after we used the covariance ratio *Cv* (relation 24) to plot the elliptic distance to the boundary.

**Figure 5:**
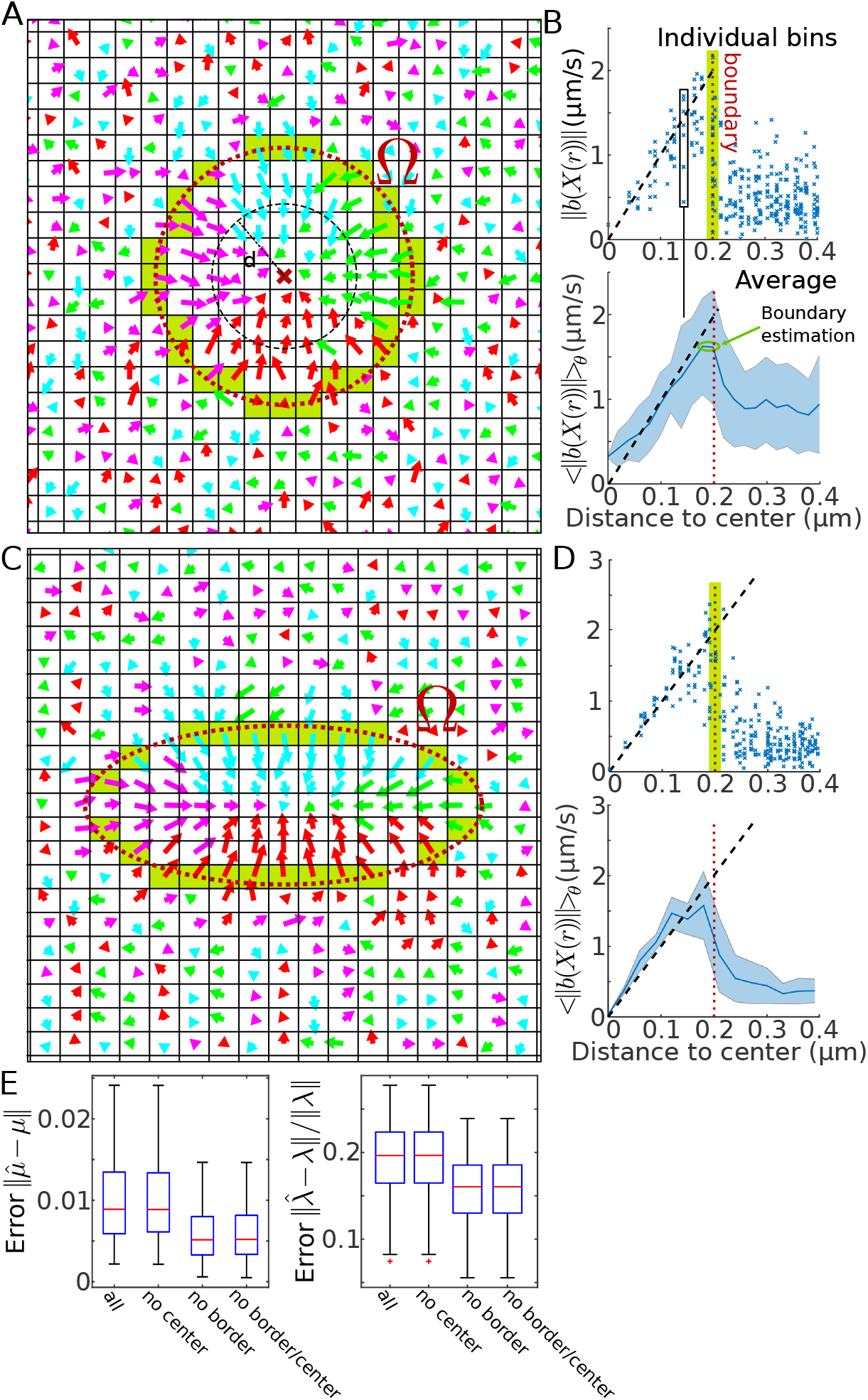
Recovery of the well depth from trajectories. **A** Vector field recovered from trajectories including the field generated by the Brownian dynamics outside the well. At the boundary of the well, there are mixed displacements (OU and Brownian), marked by the green band. **B** Upper: Drift amplitudes in each grid square versus the distance *r* to the center of the well. The expected amplitude is marked by a dashed line and the boundary with a narrow green band. Lower: Average and SD of the upper panel. **C-D** same as in A-B for the case of an ellipse where *a/b* = 2. **E** Error of the center 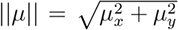 and the eigenvalue 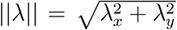 of the reconstructed ellipse in 4 cases: 1) bins falling inside the boundary are considered, 2) the center bin has been omitted 3)boundary bins are omitted and 4) when the center bin plus the ones intersecting the boundary (green bin in C) are not considered.

To evaluate the influence of bin location, we estimated the center *µ*, and eigenvalues *λ_x_* and *λ_y_* in four cases: when all bin falling inside the well, all bins except the center, all bins except the boundary and finally removing the center and the boundary bins (Fig. 5E). At this stage, we found that the best estimation is obtained by removing the center bin and the ones intersecting the boundary of the well.

## 5 Interpretation high-density regions for CaV and GPI-GFP as potential wells

The nature of high-density regions found in SPTs for various channels moving on neuronal cell membrane remains unclear. We recently reported that they could be associated with potential wells, as revealed from the voltage-gated calcium channels CaV2.1 isoform [11]. We analyze here the isoform CaV2.2 or N-type by using the three methods presented here: density (eq. (26)), least-square (equation (30)) and maximum-likelihood (see SI). We use in Fig. 6 and in table 5 only wells that contain at least 50 points from at least 5 different trajectories. All three estimators produce reasonable values of the coefficient *A* and the energy, obtained from the different methods when < 7kT. Finally, when two potential wells overlap, only one was kept. The values of the parameters are summarized in Table 5.

**Figure 6:**
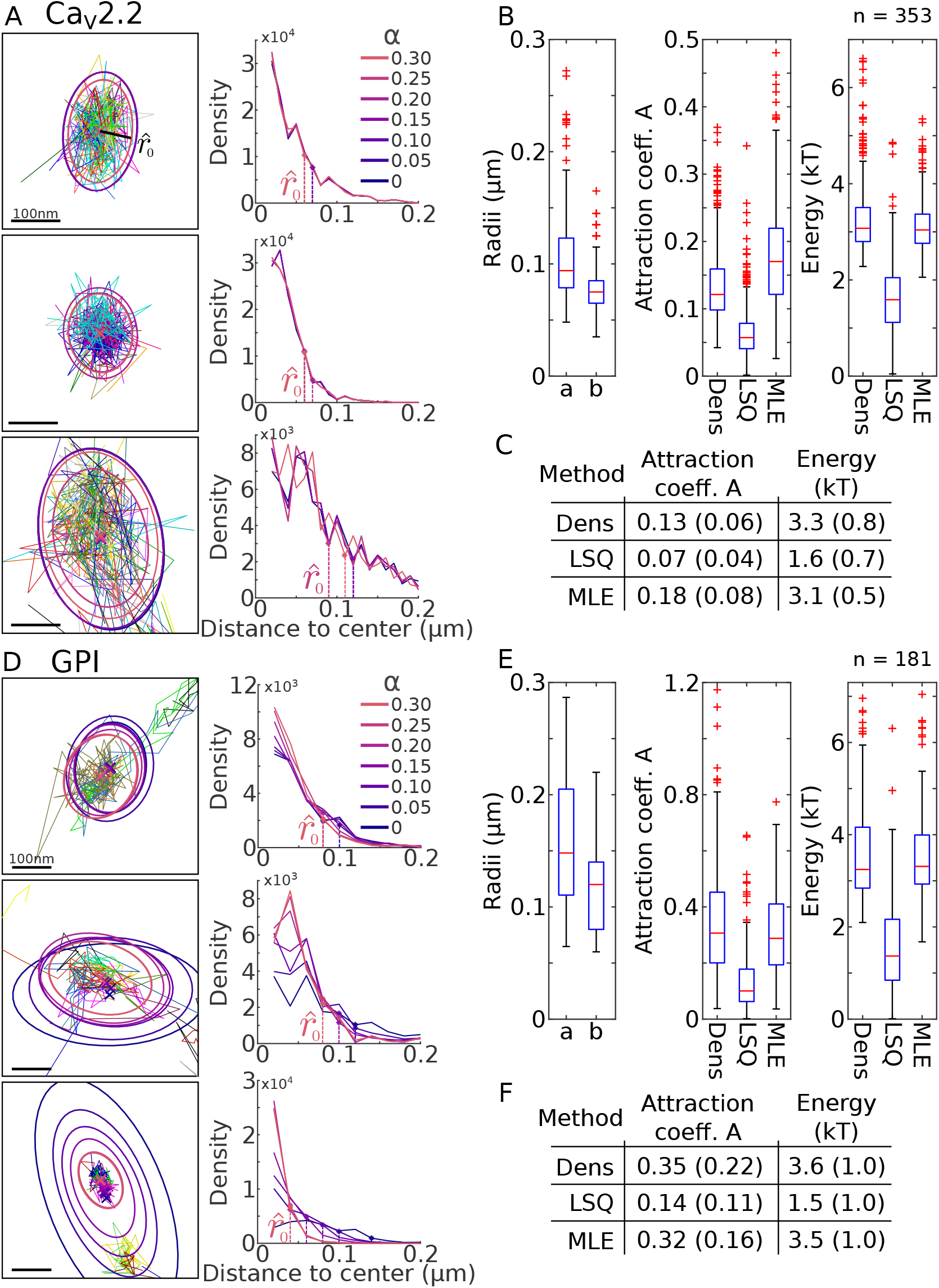
Reconstruction of wells associated to CaV2.2 and GPI. **A** 3 examples of potential wells (left) obtained from the density analysis on SPTs. The boundary of the well are estimated from various level of density *α* (right). The estimated radius 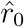 is obtained using a threshold *T* = 4% on the density. **B** Box plots for the statistics computed over 353 detected wells for the two semi-axes *a* and *b* of the ellipse, the coefficient *A* and the energy (in kT). Results are obtained for the Density, LSQ and MLE methods. **C** Summary of Mean±SD for the coefficient *A* and the energy. **D-F** same as in A-C obtained for GPI, analyzed over 181 high density regions.

We report (Fig. 6A-C) that the high density regions can be characterized as potential wells with the following characteristics: the two main axes are *a* = 90nm, *b* = 75*nm* associated with a mean energy of 3.3*kT* estimated for the density method, showing some differences with the CaV2.1 isoform [11]. Note that the distribution of energy varied with the statistical method (Fig. 6C) as we reported *E* = 3.1 ± 0.5 for the MLE and *E* = 1.6 ± 0.7 for the LSQ. At this stage, we think that the density method is more robust. To conclude, this statistical analysis suggests that to trap calcium channels, specific long-range molecular mechanisms are present in the active zone of the pre-synaptic terminal, probably associated to vesicular release molecules such as synaptotagmin. These sites retain channels for a time long enough to increase the probability of vesicular release.

We also apply our statistical methods to the case of GPI-anchored fluo-rophores (GFP) integrated into the membrane (Fig. 6D-F), which are considered to be non-interactive molecules. However, we found many regions (N=181) of high densities, which are characterized by potential wells. The elliptic axes are *a* = 140*nm* and *b* = 120*nm*, associated with an energy of *E* = 3.6, 1.5 and 3.5 for the density, LSQ and MLE methods. To conclude, although it is surprising to detect region of high densities for GPI, we found here that they can be characterized as potential wells. Possibly they correspond to regions where lipids are removed or inserted. The exact nature of these regions remain unclear and should be further investigated.

**Table 1:** Parameters used for CaV and GIP analysis.

| Parameter | GPI dataset | CaV dataset |
| --- | --- | --- |
| $\Delta x$ | 40 (nm) | 30 (nm) |
| $r_{min}$ | 30 (nm) | 20 (nm) |
| $r_{max}$ | 300 (nm) | 400 (nm) |
| $\Delta r$ | 20 (nm) | 10 (nm) |

## 6 Summary and Discussion

### 6.1 Two statistical methods to recover the nature of high density regions

We presented here two statistical methods to reconstruct from high-density regions revealed by single particle trajectories, the characteristics of a bounded potential well. The first method is based on the density of points, ignoring the causality between the successive points of the trajectories. The method is based on assuming that the potential well is generically a parabola with an elliptic base, where the diffusion coefficient is constant. In that case, the distribution of points is Gaussian Boltzmann distribution, solution of the associated Fokker-Planck equation. To recover the main parameters, we used Gaussian estimators and develop a novel strategy to recover the location of the boundary, predicted by the differences of the distribution of points: inside the well the distribution is Gaussian and outside it is uniform. We compared also the result to the classical MLE (see figure SI). The second method is based on estimating the vector field distribution at a given bin resolution ∆*x*. We use an optimal estimator to recover the characteristic of the field and the boundary is found at the discontinuity between the field computed from the energy (Ornstein-Ulhenbeck) and the fluctuations of the drift for pure Brownian motion.

At this stage, the two methods are complementary and provide certain advantages compared to classical MLE, PCA: in all cases, the center could be retrieved from simulations. However, the presence of two close wells in the automatic detection (see section 2.6) could lead to an erroneous center, which should be discarded. However, obtaining good estimtions for the gaussian parameters were clearing dependent on the methods. In particular, changing the time ∆*t* and the spatial ∆*x* steps influenced the recovery process as shown in Figs. 3 and 5. The advantage of the first (density) method based on the distribution is that we do not need to introduce an artificial grid of size ∆*x*, which could be a serious constraint in the second (drift) method because the size of the bin defines the resolution to recover the well and its boundary.

### 6.2 Nature of high density regions in empirical SPT data

Not all high density regions revealed by SPTS are due physical forces and potential wells[3]. However, when it is the case, we can recover the geometry of the well (center, curvature and boundary) and the stochastic dynamics. We applied the two methods to two types of empirical data: first, the CaV channels that mediate vesicular release at neuronal synapses and GPI-anchored control molecules that move on the surface membrane of cells. We found that high density regions for CaV have mean axes *a* = 90 and *b* = 75nm and an energy of 3.3*kT* (density method), while in principle we were expecting no regions of high density for GPI, we found some of them. There are however much less wells than CaV associated mean axes *a* = 90 and *b* = 75nm and energy 3.6*kT* (Fig. 6). Possibly the high energy of GPI well is due to a small amount of trajectories in the well, leading to a large variance.

Although the interpretation of high density regions as potential wells for AMPA receptors was first anticipated in [24] and discovered in [3], the nature of these wells and others in general, remains unclear [13]. Potential wells were found for membrane proteins such CaV [11], GAG [12, 25] and recently for G-protein [26]. Potential wells could be generated by protein clusters, or by membrane cusp of vesicle fusion points, by membrane-membrane contact at location of organelle interactions [27]. In general, potential wells are characterized long-range interaction, of the order of hundreds of nanometers. These forces could be mediated by membrane curvature, local molecular organization leading to or a combination of both.

The role of wells could vary: they retain receptors for hundreds of milliseconds to seconds at specific locations in order to increase the probability of an optimal signal transduction, such as during synaptic transmission. Transient wells allow to trap proteins to create aggregates as proposed for capsid assembly [12, 25]: once the energy of the well decreases, molecules are not interacting with the well. Another possible role is that wells could regulate the fluxes of receptors in micro-compartments such as dendritic spines [16]. They can also trap proteins in the endoplasmic reticulum [11].

Finally, it is usually difficult to identify structures hidden in SPTs. However, flows along spine neck have also been detected [3] and recently in the tubule of the endoplasmic reticulum [11]. Correlating undefined membrane geometry with an energy landscape remains difficult, because a physical model is needed to interpret them. Thus, the dynamics of receptors outside potential wells that deviates from Brownian motion is still challenging to comprehend.

